## supporting information for "Mechanism of host cell invasion by *Leishmania* through KMP-11 mediated cholesterol-transport and membrane phase transition"

### Expanded View Figures

**Evaluation of critical P/L ratio for phase transition of MΦ membrane during LD infection:**

It has been found that at P/L ratio> 0.004, DPPC remains as solution (Fluid phase) phase at physiological temperature (37°C) which resembles the actual infection condition.

Now we want to quantify the number of KMP-11 copy numbers essential for inducing the phase state change of host macrophage membranes at physiological condition.

It is known that lipid head group area[1] = 63 $Å^2$

Our as prepared Vesicles (SUVs) are of radius 70 nm.

Then area of vesicles = $4\pi r^2$ = 61544 $nm^2$.

Vesicles consist of bilayers. Assuming layer similar (layer approximation), the overall area would be= 61544 X 2= 123088 $nm^2$.

Then, no. of lipid molecules (fatty acids) in a vesicle = 123088 $nm^2$/63 $Å^2$ = 195378

Now, the working concentration of DPPC lipid = 1 mM.

So, no. of vesicles in 1 mM = 6.023 $X10^{20}$/195378 =3.08 $X10^{15}$

The number of vesicles in our working solution = 3.08 $X10^{15}$

On the other hand, protein concentration = 4 µM (since, we have taken P/L = 0.004 and in this concentration phase transition occurred at below physiological temperature).

Now, 4 µMof KMP-11 is equivalent to $6.023X10^{23}$. 4 X $10^{-6}$ = 2.4 X $10^{18}$ number of KMP-11

Therefore, in working (favorable for infection through phase transition) condition,

No. of proteins needed/ lipid vesicles = 2.4 $X10^{18}$/3.08 $X10^{15}$= 800

Considering the average size of macrophages = 5 µm which will consist of 1.25 X $10^{7}$ fatty acids. Now, 195378 fatty acids correspond to 800 number of KMP-11

So, 1.25 X $10^{7}$fatty acids correspond to [1.25 X $10^{7}$ / 195378].800.

~ 5 X $10^{4}$ copy number of KMP-11.

According to the reports, the expressed copy number of KMP-11/ LD parasite surface is.

~ 2 X $10^6$.

So, considerable amount of KMP-11 is available for the interaction and inducing infection.

Mathematical exposition of '***Hydrophobic moment and sequence symmetry-oriented phase transition model***':

Protein–lipid interactions are driven by several factors. Among these, hydrophobic interaction is the principal driving force. Although it is generally accepted that globular proteins fold with a hydrophobic core and hydrophilic exterior, the centroid of the spatial distribution of amino acid residue provides the origin of moment expansion. Spatial distribution has been described by David Silverman as a two component spherical model where interacting partners had been considered as spheres[2]. Subsequently, we constructed a '***Hydrophobic moment and sequence symmetry oriented phase transition model***', in which the nature of a helical stretch was theoretically related to the phase melting temperature of phospholipids.

Assuming the density of hydrophobicity is ρ(R) and the radial distance from the center of the sphere is R, it can be written as:

$\rho(R) = \alpha R^{\eta} - \beta R^{\mu}$,-------------------------(1)

Where, hydrophobic component will contribute an amount $4\pi\alpha R^{\eta}R^2dR$ in a shell of width dR, while the contribution of the polar component is $-4\pi\beta R^{\mu}R^2dR$.

Assuming the contribution of polar component is significantly less compared to the hydrophobic component, we can write equation (1) as

$R= [\rho(R)/\alpha]^{1/\eta}$ -----------------------(2)

Jacob Israelachvili and Richard Pashley have shown that a pair interaction free energy (ΔG) abides by the following relation[3]:

$\Delta G_H = -CRDe^{-D/D_0}$--------------------(3)

Here, hydrophobic interaction is a two-component system. Therefore, for a two component protein –lipid system, $R = {R_1 R_2}/{(R_1 + R_2)}$;where $R_1$(Peptide) and $R_2$(Lipid) are the radii of the interacting partners (Figure EV10Ei,ii). Now equation (3) will be, $\Delta G_H = -C\left({R_1 R_2}/{R_1 + R_2}\right)De^{-\frac{D}{D_0}}$

$$= -C\left({}^{1}\!/_{(1/R_1 + 1/R_2)}\right) De^{-\frac{D}{D_0}} \text{----------(4)}$$

Since the lipid component is fixed and the second component (membrane interacting domain of the protein; different peptides) is variable, we can rearrange the equation (4) as

$$\Delta G_H = -CR_1 De^{-\frac{D}{D_0}}$$

$$= -CDe^{-\frac{D}{D_0}}\left(\frac{\rho(R_1)}{\alpha}\right)^{\frac{1}{\eta}}$$

$$= -CDe^{-\frac{D}{D_0}}\left(\frac{H}{\alpha A}\right)^{\frac{1}{\eta}} \text{------------------------- (5)}$$

Where, H is the net hydrophobicity and A is the area of hydrophobic face.

From our experimental and theoretical calculations, it has been established that protein-lipid interaction and phase transition is dependent on the hydrophobic moment of the proteins stretch, therefore, equation (4) can be represented as

$$\Delta G_H = -CDe^{-\frac{D}{D_0}}\left(\frac{\mu_H}{\alpha A}\right)^{\frac{1}{\eta}} \text{------------------------------------------- (6)}$$

Using the mirror sequence constituting residue number (N) and different combinations of polar (P) and hydrophobic (H) residues of KMP-11 amino–terminal stretch (1-19) (Table EV5), we showed the linear correlation between the hydrophobic moment, $\mu_H$ and N. The linear equation is expressed (Figure EV10F) as:

$$\mu_H = 0.018N + 0.227 \text{----------------------------(7)}$$

Now, combining equation (6) and (7), we get

$$\Delta G_H = -CDe^{-\frac{D}{D_0}}\left(\frac{0.018N+0.227}{\alpha A}\right)^{\frac{1}{\eta}} \text{------------------------ (8)}$$

Since hydrophobic free energy is the strongest component for the phase transition to occur, we can write, $\Delta G_H = \Delta G_T$; where $\Delta G_T$ is the free energy change during gel/fluid phase transition. Now, Thermodynamic equation of free energy would be,

$\Delta G_T = \Delta H_T - T_T \Delta S_T$ -------------------------------- (9) where, $T_T$ stands for the phase transition temperature of phospholipid.

Combining equation (8) and (9), we can write,

$$T_T \Delta S_T = \Delta H_T + CDe^{-\frac{D}{D_0}} \left(\frac{0.018N + 0.227}{\alpha A}\right)^{\frac{1}{\eta}}$$

Or, $T_T = \Delta H_T / \Delta S_T + \frac{CDe^{-\frac{D}{D_0}} \left(\frac{0.018N + 0.227}{\alpha A}\right)^{\frac{1}{\eta}}}{\Delta S_T}$ --------------------------------- (10)

Assuming $\eta = 1$, we can write equation (10) as

$T_T = \Delta H_T / \Delta S_T + \frac{K(0.018N + 0.227)}{\Delta S_T}$ ------------------------------------- (11)

Where, $K = \frac{CDe^{-\frac{D}{D_0}}}{\alpha A}$

Linear fitting (Figure EV10G) of the experimental data of transition temperature ($T_m$) against N has given us simplified linear correlation e.g

$T_m = 42.02 - 0.466N$---------------------------------------------- (12)

The proposed model indicated a strong correlation between the sequence amino acid distribution and the protein mediated phase transition temperature of phospholipid membranes.

**Table EV1: Fluorescence lifetimes($T_{avg}$,nsec) of DiD labeled DPPC SUVs in presence and absence of r-KMP-11 at protein/lipid 0.02(molar ratio). The data is represented as Mean±SD evaluated from three independent experiments.**

| Systems | $T_1$ | $T_2$ | $T_{avg}$ | $X^2$ |
|---|---|---|---|---|
| **DiD - DPPC** | 3.397±0.121 | 5.695±0.152 | 4.107±0.115 | 1.091 |
| **DiD-DPPC-r-KMP-11** | 2.813±0.098 | 4.438±0.132 | 3.968±0.109 | 1.049 |

**Table EV2: Binding constants ($K_a$), cooperative index ($n$) and Stern-Volmer constants ($K_{sv}$) of different protein mutants with DPPC model membranes.Binding experiments were performed with DiD labeled DPPC SUVs in presence r-KMP-11.**

| Systems | $K_a$ ($M^{-1}$) | $n$ | $K_{sv}$($M^{-1}$) |
|---|---|---|---|
| r-KMP-11-DPPC | $3.504 \times 10^6$ | 1.450 | ------ |
| Y5W | -------- | ------- | 4.909 |
| Y5W-DPPC | $5.154 \times 10^7$ | 2.117 | 2.453 |
| F19W | ---------- | ----------- | 4.665 |
| F19W-DPPC | $4.351 \times 10^7$ | 1.822 | 3.146 |
| F30W | ---------- | -------- | 5.142 |
| F30W-DPPC | $2.200 \times 10^7$ | 1.772 | 4.716 |
| Y48W | ----------- | -------- | 1.799 |
| Y48W-DPPC | $8.508 \times 10^6$ | 1.945 | 1.455 |
| F51W | --------- | --------- | 4.268 |
| F51W-DPPC | $1.540 \times 10^7$ | 1.682 | 4.103 |
| F62W | ------- | ------- | 3.177 |
| F62W-DPPC | $2.674 \times 10^5$ | 1.562 | 4.454 |
| F77W | -------- | -------- | 9.862 |
| F77W-DPPC | $3.000 \times 10^6$ | 1.624 | 9.567 |
| Y89W | -------- | --------- | 4.454 |
| Y89W-DPPC | $6.205 \times 10^6$ | 2.364 | 4.152 |

**Table EV3: Chain melting temperature of DPPC membrane using different r-KMP-11 protein (P)/DPPC Lipid(L) molar ratio as monitored by measuring the polarsiation of Laurdan and REES of NBD PC.**

| P/L ratio | Chain melting temperature using laurdan | Chain melting temperature using NBDPC |
|---|---|---|
| 0 | 44.8 | 44.4 |
| 0.0002 | 47.4 | 46.2 |
| 0.001 | 48.0 | 43.9 |
| 0.002 | 40.4 | 36.5 |
| 0.0025 | 39.5 | 37.7 |
| 0.004 | 38.7 | 33.9 |
| 0.02 | 33.5 | 33.2 |

**Table EV4: Efficiency of energy transfer(E) and donor-acceptor distances ($r_0$) of different tryptophan residues of eight single tryptophan mutants. Here, tryptophan was the donor and DHE in DPPC SUVs was used as the acceptor molecules. Here, Binding of different mutants with DPPC SUVs was examined to determine the orientation of KMP-11 in DPPC membrane.**

| Mutants | E | $r_0$ (Å) |
|---|---|---|
| Y5W | 0.20 | 11.4 |
| F19W | 0.10 | 12.8 |
| F30W | 0.08 | 18.2 |
| Y48W | 0.06 | 14.1 |
| F51W | 0.02 | 15.6 |
| F62W | 0.01 | 24.7 |
| F77W | 0.06 | 15.8 |
| Y89W | 0.08 | 13.6 |

**Table EV5: Different combinations of the amino acid residues of the amino-terminal stretch (1-19AA)of KMP-11 based on their polarity and hydrophobicity. Here, N denotes the number of amino acid residues and $\mu_H$ is the hydrophobic moments calculated by the Helical wheel model. The membrane embedded residues and the free energy changes were calculated in OPM server.**

| Serial No. | Peptide sequences | N | μH | Embedded residues | ΔG(Kcal/mole) |
|---|---|---|---|---|---|
| 1 | MATTYEEFSAKLDRLDQEF<br>HHPPHPPHPHPHPPHPPPH | 15 | 0.505 | 1,5,8,12,15,19 | -9.7 |
| 2 | MATTYEEFSAKLDRLFDQE<br>HHPPHPPHPHPHPPHHPPP | 11 | 0.453 | 1,2,5,8,12 | -7.7 |
| 3 | MATTYEEFSAKLDLRDQEF<br>HHPPHPPHPHPHPPPPHPH | 9 | 0.394 | 1,5,8,12,19 | -8.4 |
| 4 | MATTYEEFSAKLDLRDQEF<br>HHPPHPPHPHPHPHPPPPH | 7 | 0.354 | 1-5,8 | -7.3 |
| 5 | MATTYEEFSAKLLDRDFQE<br>HHPPHPPHPHPHHPPPHPP | 5 | 0.306 | 1-6,8 | -6.8 |
| 6 | MATTYEEFSAKDLRDLFQE<br>HHPPHPPHPHPPHPPHHPP | 3 | 0.292 | 1-2,4-5,8 | -6.6 |

**Table EV6: Disease causing proteins with their membrane binding domains and respective N values.**

| DISEASE | PROTEIN | MEMBRANE BINDING STRETCH | N |
|---|---|---|---|
| Whooping cough | Adenylcyclase toxin of B. pertusis | **LFGRAPEVIARA [HHHHHHPHHHHH]** | 11 |
| Diarrhoeal diseases and deep wound infections | Aerolysin | **RLFSLGQGVCGDK [HHHPHHPHHPHPP]** | 9 |
| Gas gangrene | Alpha toxin from C. perfringens | **KDNSWYLAYSIPDTGES [PPPPHHHHHPHHPPHPP]** | 7 |
| Anthrax | Anthrolysin O | **KVSIGGTTLYPTATISH [PHPHHHPPHHHPHHHPP]** | 11 |
| Food poisoning/gastrointestinal illnesses | Clostridium perfringens enterotoxin | **QSLGDGVKDHYVDISL [PPHHPHHPPPHHPHPH]** | 11 |
| Cholera | Colicin A | **EVESWVLSGIASSVAL [PHPPHHHPHHHPPHHH]** | 13 |
| Respiratory tract infection | Delta toxin | **IRKEENGNVTIITQNNKQ [HHPPPPHPHPHHPPPPPP]** | 5 |
| Gas gangrene, | Epsilon toxin | **TTTHTVGTSIQATAKFTVPFNETGVSLTTSYSFANTN [PPPPPHHPPHPHPHPHPHHHPPPHHPHPPPHPHHPPP]** | 7 |
| Cardiotoxic effects | Equinatoxin II | **QKDRGPVATGAVGVLAYLMSD [PPPHHHHHPHHHHHHHHHHPP]** | 11 |
| Hemolytic activity | Gama hemolysin | **SRTTYSDLIKRMIWPF [PHPPHPPHHPHHHHHH]** | 5 |
| endophthalmitis | Hemolysin BL | **LSEIEQTNNGDTAL [HPPHPPPPPHPPHH]** | 13 |
| Hemolytic activity | Hemolysinlectin | **NVLATQTLENTSSQTQE [PHHHPPPHPPPPPPPPP]** | 9 |
| Forms cytotoxic pores in CD59-positive cells, promotes adherence to and invasion of human liver cells | Intermedilysin | **ATGLAWEPWRLIYS [HPHHHHPHHHHHHP]** | 9 |
| Forms cytotoxic pores in CD59-positive cells, | Lectinolysin | **EVFRSATNIG [PHHHPHPPHH]** | 5 |

| | | | |
|---|---|---|---|
| promotes adherence to and invasion of human liver cells | | | |
| Kill leucocytes | Leucocidin F | **RWNGFYWAGANY [HHPHHHHHHHPH]** | 11 |
| listeriosis | Listeriolysin O | **ECTGLAWEWWRTVID [PPPHHHHPHHHPHHP]** | 7 |
| | Perfringolysin O | **YDVPLTNNINVSIWGTTLYPGSST [HPHHHPPPHPHPHHHPPHHHHPPP]** | 5 |
| | Sticholysin II | **SVPFDYNWYSNWWDVK [PHHHPHPHHPPHHPHP]** | 7 |
| Disrupts cytoplasmic integrity of erythrocytes, leukocytes, macrophages, platelets, epithelial cells | Streptolysin O | **TGLAWEWWRKV [PHHHHPHHHPH]** | 7 |
| Cytotoxic to endothelial cells, epithelial cells, macrophages, and neutrophils | Suilysin | **CTGLAWEWWRTVY [PPHHHHPHHHPHH]** | 7 |
| Cholera | Vibrio choleraecytolysin | **AYKHYYVVGAHQSYH [HHPPHHHHHHPPPHP]** | 5 |
| Cholera | Vibrio vulnificushemolysin | **MAHVTLQSLSNN [HHPHPHPPHPPP]** | 5 |
| Severe Acute Respiratory Syndrome | Coronavirus Protein 6 | **MFHLVDFQVTIAEILIIIIMRTFRIAIWNLDVIISSIVR [HHPHHPHPHPHPPHHHHHHHPPHPHHHHPHP]** | 17 |
| Hepatitis C | HCV NS5A (Hepatitis C virus) | **DTSWLRDVWDWVCTVLSDFRVWLQAKLL [PPPHHPPHHPHHPPHHPPHPHHHPPPHH]** | 17 |
| Diptheria | Diptheria toxin | **SLTGTNPVFAGANYAAWAVN [PHPPPPHHHHPHPHHHHHHP]** | 11 |
| E. Coli | Cytolysin A | **KTVEVVKNAIETADGLDLYNKYLD [PPHPHHPPHHPPHPPHPHHPPHHP]** | 7 |

**Table EV7: Primers of the single tryptophan mutants of KMP-11**

| Tryptophan Mutations | Primers (5' → 3') |
|---|---|
| Y5W | Fwd: CCATGGCCACCACGTGGGAGGAGTTTTCGGCG<br>Rev: GCCGAAAACTCCTCCCACGTGGTGGCCATGG |
| Y48W | Fwd: CGAGATGAAGGAGCACTGGCAGAAGTTCGAGCGCAT<br>Rev: ATGCGCTCGAACTTCTCCCAGTGCTCCTTCATCTCG |
| F62W | Fwd: GCTCGTGCATCTTCTTGTTCCACTTCTCTGTGTGTTCCTT<br>Rev:<br>AAGGAACACACAGAGAAGTGGAACAAGAAGATGCACGAGC |
| Y89W | Fwd: AGCAGAAGGCTGCGCAGTGGCCGTCCAAGC<br>Rev: GCTTGGACGGCCACTGCGCAGCCTTCTGCT |
| F19W | Fwd: CCGCCTGGATGAGGAGTGGAACCGGAAGATGCAGG<br>Rev: CCTGCATCTTCCGGTTCCACTCCTCATCCAGGCGG |
| F30W | Fwd: GAGCAGAACGCCAAGTGGTTTGCGGACAAGCCGG<br>Rev: CCGGCTTGTCCGCAAACCACTTGGCGTTCTGCTC |
| F51W | Fwd: GGAGCACTACGAGAAGTGGGAGCGCATGATCAAGGA<br>Rev: TCCTTGATCATGCGCTCCCACTTCTCGTAGTGCTCC |
| F77W | Fwd: CGGAGCACTTCAAGCAGAAGTGGGCCGAGCTGCT<br>Rev: AGCAGCTCGGCCCACTTCTGCTTGAAGTGCTCCG |

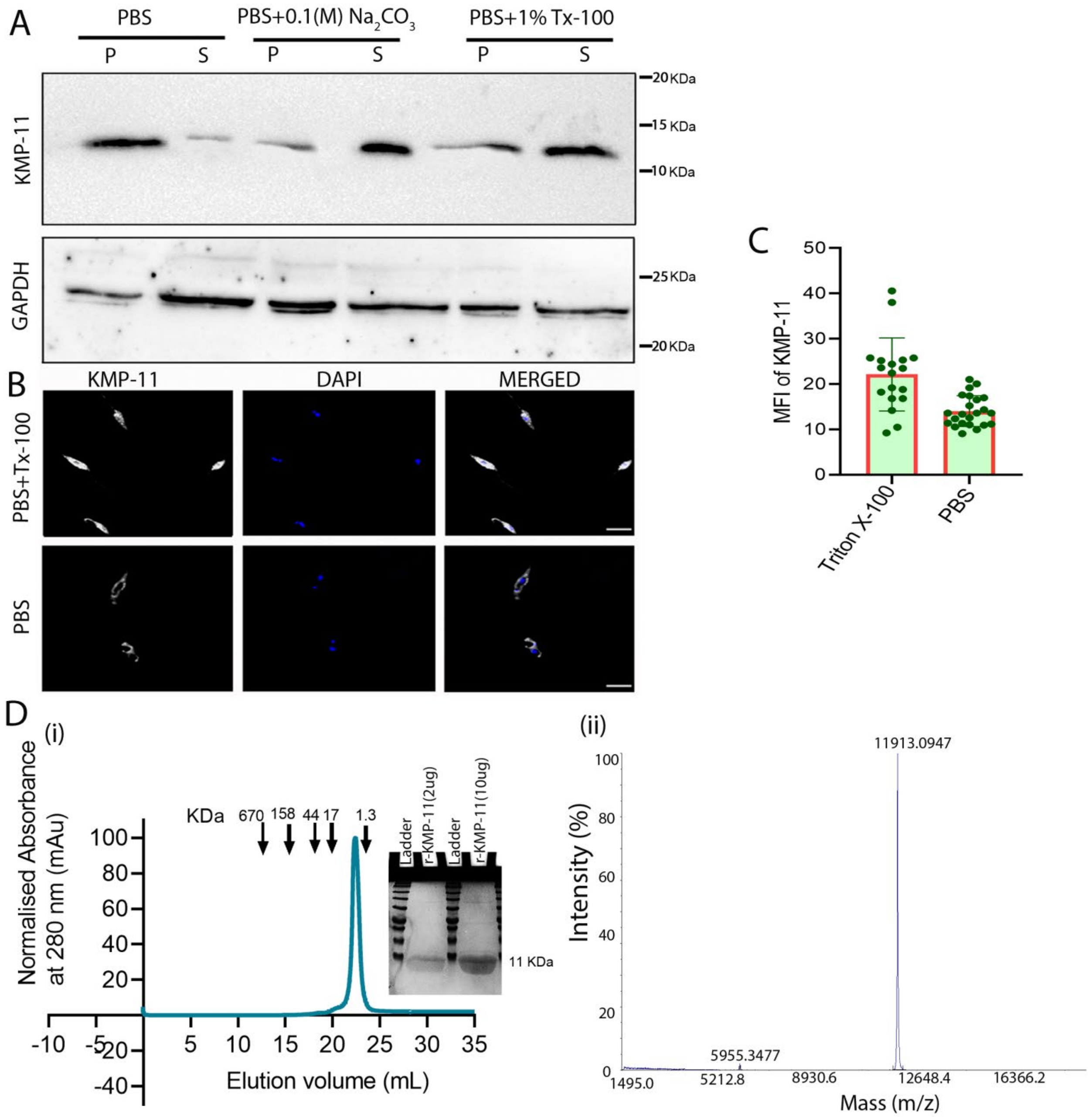


**Figure EV1: KMP-11 is exposed on LD surface**. (A) Western Blot showing level of KMP-11 in pellet (P) and soluble fraction (S) of PBS, PBS with 0.1(M) $Na_2CO_3$, and PBS with 0.1% Triton X-100. It shows that KMP-11 accumulates in soluble fractions in the presence of detergent. (B) Confocal microscopy showing KMP-11 in the surface, flagella, and cytoplasm of LD in case of permeabilization with 0.1% Triton, while in absence of permeabilization, it is retained on LD surface. Scale bar is 5µm. (C) Plot of mean fluorescence intensity (MFI) of KMP-11 on LD surface in PBS and in PBS with 0.1%TX-100. This data shows that a significant amount of KMP-11 is accessible in the LD even without permeabilization. MFI has been calculated by selecting LDs (n=17 for PBS with 0.1%TX-100 condition and n=23 for PBS condition) from the confocal microscopic images of three biological replicates for each condition. Fiji Image J was used to evaluate the intensity. (D).(i) Gel filtration chromatography. Absorbance at 280 nm was monitored continuously to identify the purified protein. Standard molecular mass markers (thyroglobulin, 670 KDa; ϒ-globulin, 158 KDa; ovalbumin, 44 KDa; myoglobin, 17 KDa; and vitamin B12, 1.35 KDa) were chromatographed on the same column (arrows) under the same conditions. The apparent molecular mass of our purified r-KMP-11 is ~11 KDa. Inset shows the SDS page of our purified r-KMP-11. Two different amounts of r-KMP-11(5 and 20 μg) were loaded in 15% SDS containing acrylamide gel. (ii) MALDI-TOF mass spectra of our purified r-KMP-11.

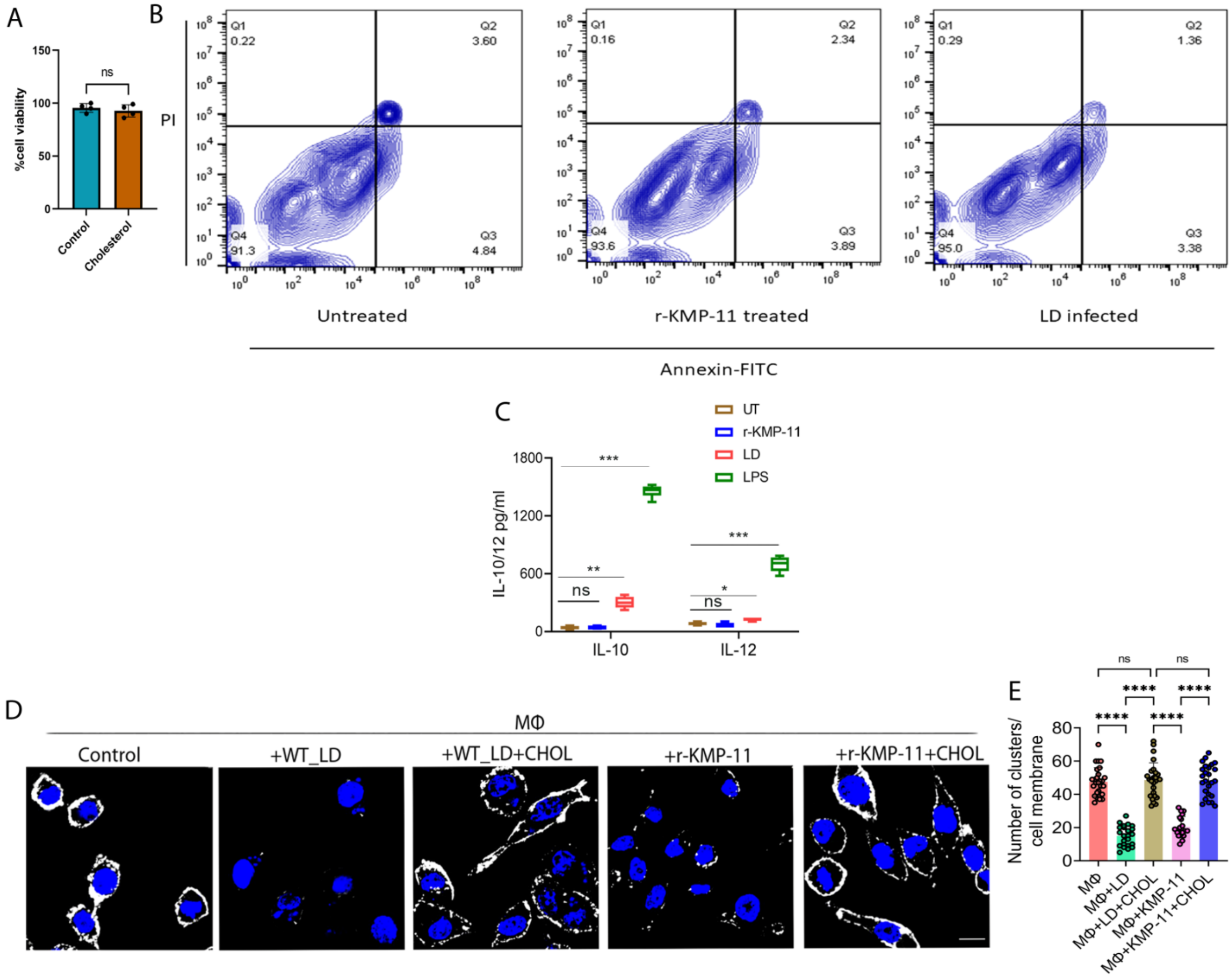


**Figure EV2: Effect of KMP-11.** (A) MTT assay shows no toxic effect of 1.5 mM cholesterol treatment on raw 264.7 MΦ cell line. Data are represented as Mean ± SD derived from three biological replicates. (B). Flow Cytometry representing the percentage of apoptotic MΦ population in response to r-KMP-11 by Annexin V-propidium iodide (PI) staining. MΦs were treated with 50μM of r-KMP-11 for 12 hrs or left untreated or infected with LD and stained with Annexin V-FITC and propidium iodide. Percentage of apoptotic cell population was measured using flow cytometry. Data represented as contour plots and double-positive cells (Annexin V-FITC and PI) demonstrate the typical apoptotic cell population. This figure is a representative of three independent experiments. (C) Level of IL10 and IL12 in MΦs infected with LD, or treated with 50μM KMP-11, or LPS (1mg/ml) for 12 hr. Data are represented as Mean ± SD derived from three biological replicates. The statistical significance of the data has been derived using ANOVA in GraphPad Prism (version 9) application. UT stands for untreated. *p<0.05,**p<0.005,***<0.001.(D).Confocal images of membrane rafts under treatment of r-KMP-11 and LD infection, and the reassembly of the raft after CHOL treatment. CTX-B-FITC was used as the raft marker. Images of control MΦ, LD-MΦ, LD- MΦ with CHOL, KMP-MΦ and KMP-MΦ with CHOL have been shown. The scale bar in each image is 10μm. In order to represent Ctxb clusters, all representative images for each experimental condition are uniformly represented with high contrast. (E). Plot of the number of punctate raft clusters per cell membrane for control MΦ, LD-MΦ, LD- MΦ with CHOL, KMP-MΦ and KMP-MΦ with CHOL. Data has been represented as Mean ± SD.MΦ cells (n=21) for each condition were collected from three independent confocal images and were evaluated using Fiji Image J software. The level of significance has been estimated using ANOVA in GraphPad Prism (version 9) application. ****p<0.0001.

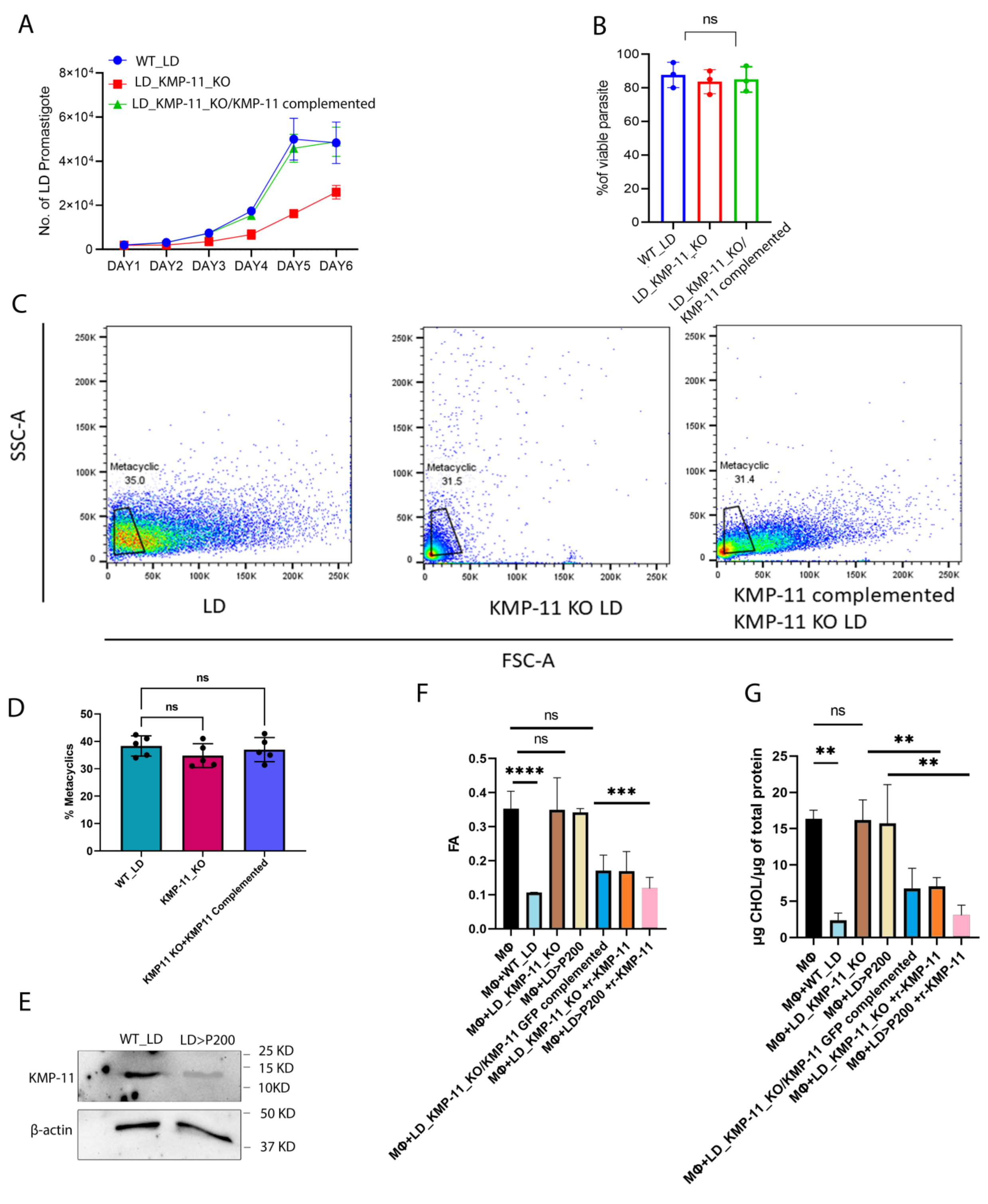


**Figure EV3: Complementation and characterization of LD_KMP-11_KO lines.** (A).*In vitro* growth kinetics comparing WT_LD,LD_KMP11_KO and LD_KMP11_KO /KMP-11 GFP complemented lines with equal inoculums ($10^2$/ml) over a period of 6 days. Growth rate of

WT_LD is significantly higher post $4^{th}$ day of *in vitro* culture.(B) MTT assay comparing viability of WT_LD,LD_KMP-11_KO and LD_KMP11_KO /KMP-11 GFP complemented lines in equal number ($10^5$) of WT_LD ,LD_KMP-11 KO and LD_KMP11_KO /KMP-11 GFP complemented lines from a $6^{th}$ day culture used for primary inoculum. Data has been presented as Mean±SD derived from three independent biological replicates. The level of significance has been estimated using ANOVA in GraphPad Prism (version 9) application.(C). Detection of stationary LD promastigotes by Flow cytometry on the basis of scatter light. First panel showing the WT_LD, middle panel showing the KMP-11 KO LD and the right panel showing the KMP-11 complemented KMP-11 KO LD. This data is a representative of three independent experiments. (D). Flow cytometry representing percent metacyclic in stationary phase ($6^{th}$ day culture) of WT_LD, LD_KMP-11_KO and LD_KMP11_KO /KMP-11 GFP complemented lines. WT_LD lines showing slightly higher percentage of metacyclic than LD_KMP-11_KO and LD_KMP11_KO /KMP-11 GFP complemented lines. Data has been represented as Mean ± SD(n=5). (E) A significant higher expression of KMP-11 was observed by western blot in case of WT_LD (passage 3, P3) as compared to LD lines having more than 200 passages (LD>P200) in M199 *in vitro* culture. Macrophage membrane fluidity change and CHOL depletion by KMP-11. (F). The values of FA of MΦ membrane upon the treatment of WT_LD, LD_KMP-11_KO, LD>P200, LD_KMP-11_KO/KMP-11 GFP Complemented and LD_KMP-11_KO, LD>P200 in presence of exogenous r-KMP-11 (50 μM). Data has been represented as Mean ± SD (n=2).The level of significance has been estimated using ANOVA in GraphPad Prism (version 9) application.****p<0.0001, ***p<0.001. (G). Membrane fraction was isolated from the MΦs by ultracentrifugation. Total membrane cholesterol (free and esterified) was estimated using the Amplex Red kit and expressed as μg of cholesterol/μg of total protein. MΦ membranes were treated with WT_LD, LD_KMP-11_KO, LD>P200, LD_KMP-11_KO/KMP-11 GFP Complemented and LD_KMP-11_KO, LD>P200 in presence of exogenous r-KMP-11 (50 μM). Data has been represented as Mean ± SD (n=2).The level of significance has been estimated using ANOVA in GraphPad Prism (version 9) application.**p<0.005.

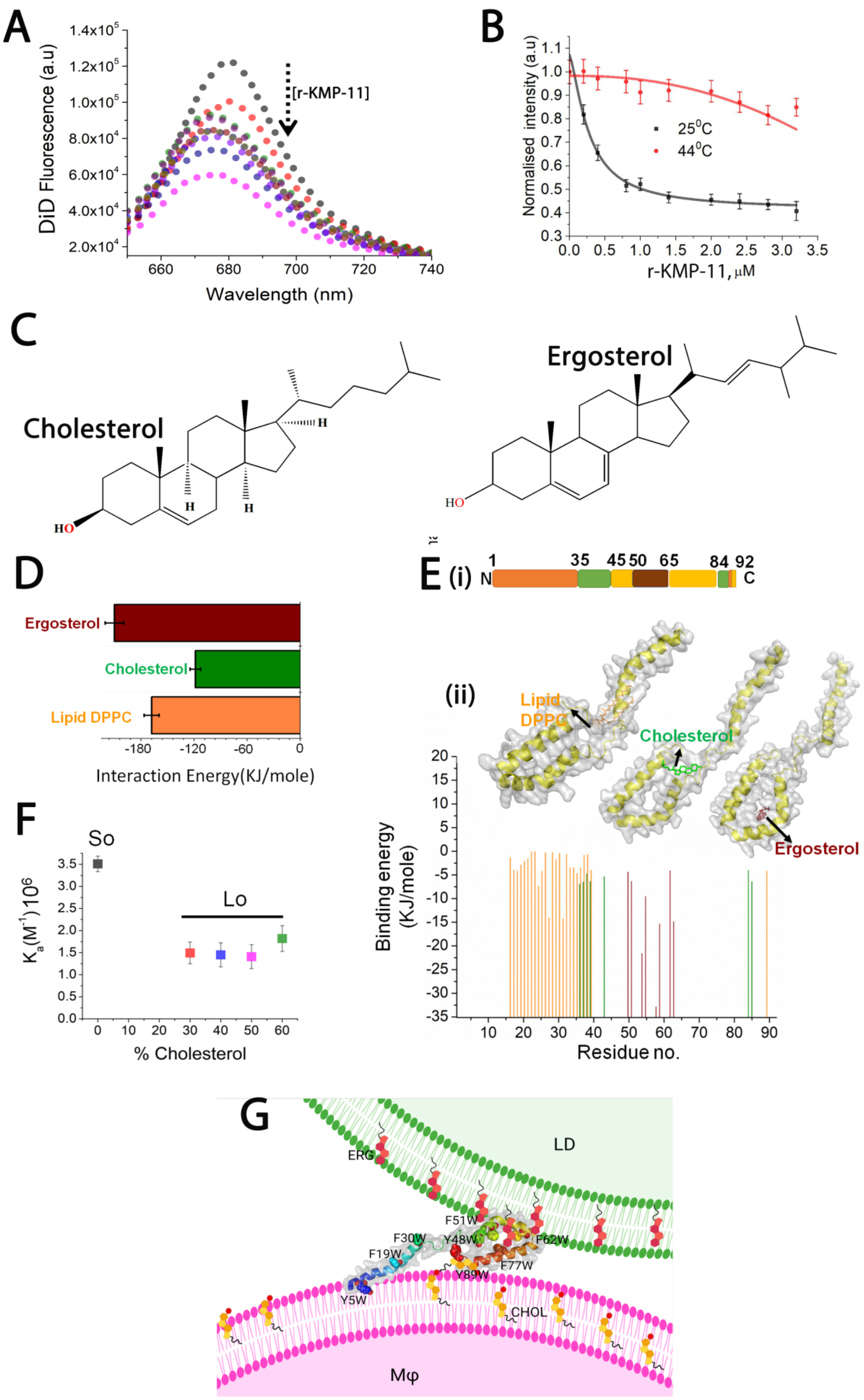
A
DiD Fluorescence (a.u)
[r-KMP-11]
Wavelength (nm)
B
Normalised intensity (a.u)
25$^0$C
44$^0$C
r-KMP-11, μM
C
Cholesterol
Ergosterol
D
Ergosterol
Cholesterol
Lipid DPPC
Interaction Energy(KJ/mole)
E (i)
N
C
(ii)
Lipid DPPC
Cholesterol
Ergosterol
Binding energy (KJ/mole)
Residue no.
F
So
Lo
$K_a(M^{-1})10^6$
% Cholesterol
G
LD
ERG
F51W
F30W
Y48W
F62W
F19W
F77W
Y89W
Y5W
CHOL
Mφ

**Figure EV4: KMP-11 binding with model DPPC SUVs.** (A). Representative DiD fluorescence titration spectra shows the binding of r-KMP-11 with DPPC model membranes as was monitored by the decrease in DiD fluorescence intensity (left y axis) as well as the blue shifts of the fluorophore (right y axis) with increasing concentration of added proteins. (B). Plot of normalized membrane embedded DiD fluorophore intensities against increasing WT r-KMP-11 concentration to estimate the binding of r-KMP-11 with DPPC at two different temperatures (25°C and 44°C). Two different temperatures were used to see the binding of r-KMP-11 with gel and fluid DPPC membrane. The solid lines indicated the fit of the data to evaluate the binding affinities. Our data showed the binding affinity with Lo/gel phase DPPC (25°C) is considerably higher than the Ld/fluid phase DPPC membrane (44°C). (C). Chemical structures of cholesterol and ergosterol. Region-specific binding of lipid and sterol molecules with KMP-11. (D). The binding energy variation of KMP-11 with DPPC lipid, CHOL and ERG as obtained by docking analysis. (E). A schematic drawing of the overall protein sequence which is marked based on the binding locations of CHOL (green), ERG (brown) and DPPC lipid(orange) as obtained from the molecular docking study. The best posed docked structure of KMP-11 as obtained from docking study was shown. The amino-terminal domain shows the DPPC lipid binding domain, 35-45 AA region and 85-90 AA regions are found to be CHOL binding region,50-70 AA region is the ERG binding motif. Plot of binding energy of individual lipid and sterol components against the residue number of KMP-11 as obtained from gemdock tool has been shown. Snapshots of the best posed docked structures for CHOL-KMP-11; ERG-KMP-11 and DPPC lipid-KMP-11 as obtained from the molecular docking study. (F). Binding of r-KMP-11 with DPPC SUVs containing different mole percentages of CHOL as obtained from the membrane embedded DiD quenching assay. Data indicated the notable decrease in the r-KMP-11 binding with DPPC SUVs containing different percentage CHOL. Here, So and Lo stand for solid order and liquid order states respectively. This data signifies higher binding affinity of r-KMP-11 towards So domain in comparison to Lo domain. (G). The plausible orientation of KMP-11 during attachment of the parasite (DPPC-ERG) with MΦ (DPPC-CHOL) membrane.

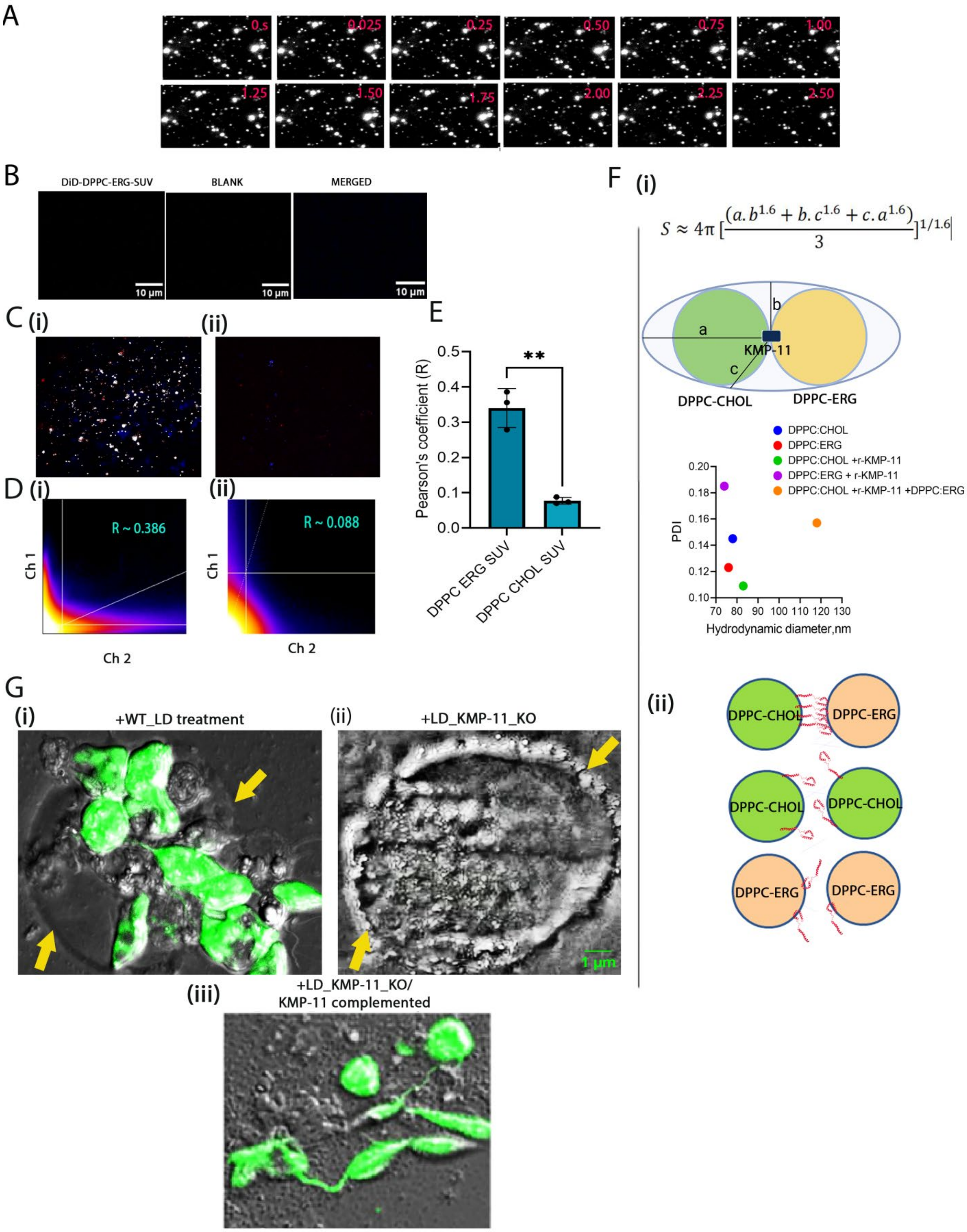
A
0 s
0.025
0.25
0.50
0.75
1.00
1.25
1.50
1.75
2.00
2.25
2.50
B
DiD-DPPC-ERG-SUV
BLANK
MERGED
10 µm
C (i)
(ii)
D (i)
(ii)
R ~ 0.386
R ~ 0.088
Ch 1
Ch 2
E
Pearson's coefficient (R)
**
DPPC ERG SUV
DPPC CHOL SUV
F (i)
$S \approx 4\pi \left[\frac{(a.b^{1.6} + b.c^{1.6} + c.a^{1.6})}{3}\right]^{1/1.6}$
a
b
c
KMP-11
DPPC-CHOL
DPPC-ERG
DPPC:CHOL
DPPC:ERG
DPPC:CHOL +r-KMP-11
DPPC:ERG + r-KMP-11
DPPC:CHOL +r-KMP-11 +DPPC:ERG
PDI
Hydrodynamic diameter,nm
(ii)
DPPC-CHOL
DPPC-ERG
G
(i) +WT_LD treatment
(ii) +LD_KMP-11_KO
1 µm
(iii) +LD_KMP-11_KO/
KMP-11 complemented

**Figure EV5: The bridging nature of KMP-11.** Colocalization study on PEG cushioned SLB. (A). Time lapse images of the DiD labeled DPPC: ERG SUVs on DPPC: CHOL SLB platform in presence of r-KMP-11. The dimension of each image was kept 32.19x25.94 $\mu m^2$. (B). TIRF microscopic images of DiD labeled DPPC ERG SUVs (red channel), blank (blue channel) and the merged image. (C). (i) Colocalised micrograph of DiD labeled DPPC ERG SUVs (red channel) and Alexa-488 maleimide labeled r-KMP-11 (blue channel). (ii) colocalised micrograph of DiD labeled DPPC CHOL SUVs (red channel) and Alexa-488 maleimide labeled r-KMP-11 (blue channel). (D). Scatter plots of colocalisation of (i) DiD labeled DPPC ERG SUVs (red channel) and Alexa-488 maleimide labeled r-KMP-11 (blue channel) system and (ii) DiD labeled DPPC CHOL SUVs (red channel) and Alexa-488 maleimide labeled r-KMP-11 (blue channel). The R (pearson's coefficient) values stand for the degree of colocalisation. (E).Plot shows the values of pearson's coefficients obtained from TIRF imaging using DPPC:CHOL and DPPC: ERG SUVs. Data were presented as Mean±SD derived from three independent biological replicates. The level of significance has been estimated using ANOVA in GraphPad Prism (version 9) application. $^{**}p<0.005$. (F). (i)Dynamic light scattering (DLS) study to estimate the hydrodynamic radii of different sterol-containing SUVs. We observed comparable hydrodynamic radii for DPPC-ERG and DPPC-CHOL. Hydrodynamic radius increased significantly when we added r-KMP-11 to the equimolar mixture of DPPC-ERG and DPPC-CHOL. Implying ellipsoid approximation while two vesicles are attached by a KMP-11 bridge, we calculated the average surface area of the combined system, which was found to be ~ 43870.42 $nm^2$. Interestingly, we found that our calculated surface area matched well with the surface area obtained from DLS data for DPPC-ERG + DPPC-CHOL+r-KMP-11 system (observed surface area ~41094 nm2), which further suggests that KMP-11 can act as a bridging molecule between CHOL and ERG rich membranes. Here, PDI stands for poly-dispersity index. (ii). Different orientations of KMP-11 in SUV environment in presence of CHOL and ERG in SUVs. (G). Images representing interaction of CFSE labeled WT_LD and LD_KMP-11_KO and complemented LD lines on supported Lipid Bilayer (SLB) composed of DPPC CHOL 30%. Significant attachment of (i) WT_LD parasites was observed on lipid surface which is absent for (ii) LD_KMP-11_KO parasites. Attachment is restored for (iii) LD_KMP-11_KO/complemented parasites on SLB surface. The yellow arrows indicate the edge of the supported lipid bilayer.

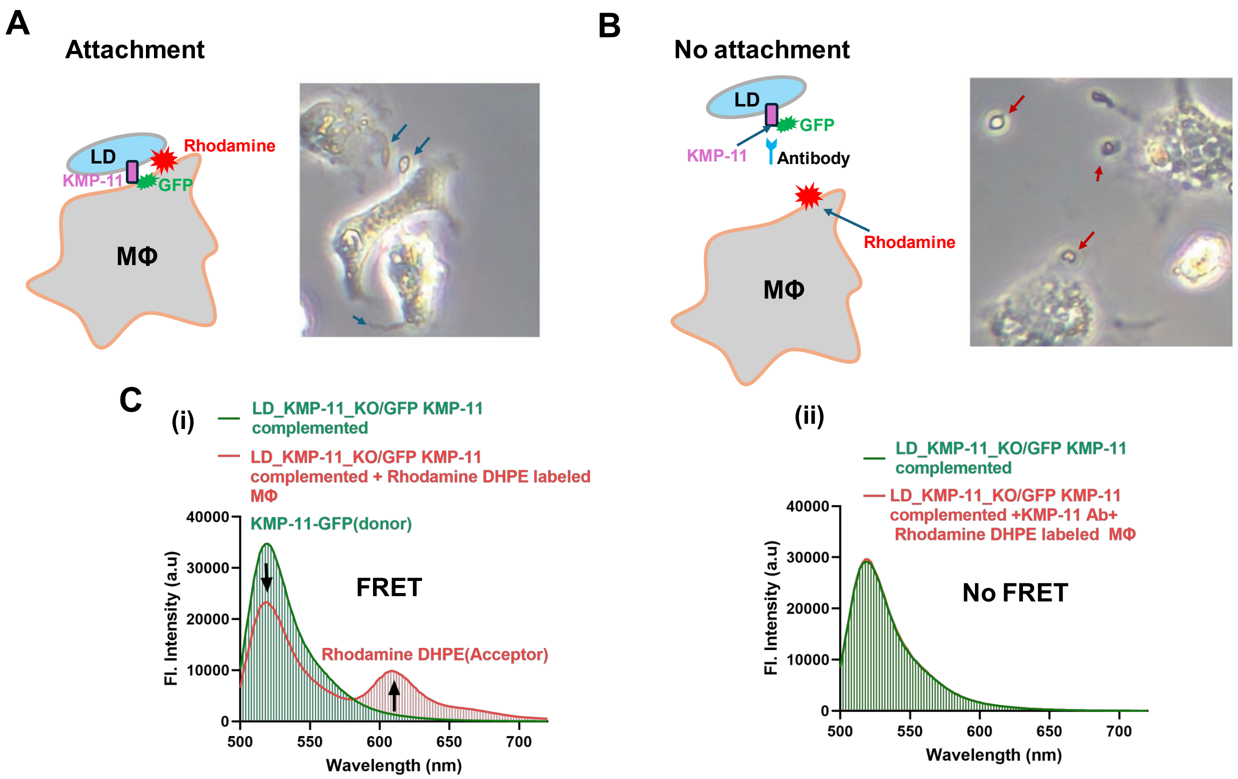


**Figure EV6: Model of FRET experiment to determine the binding of KMP-11 on LD with Macrophage membrane during attachment.** (A.) **Condition 1:** GFP tagged KMP-11 in the KMP-11 complemented LD lines was used as donor and Rhodamine DHPE in macrophage membrane were used as acceptor. In attached condition, we can see significant FRET if donor and acceptor molecule come close to each other. Bright field image shows the attached LD (blue arrow marked) with MΦ. (B). **Condition 2:** We incubated the GFP tagged KMP-11 in the KMP-11 complemented LD lines with anti KMP-11 antibody to reduce the attachment. In unattached condition, there should not occur FRET as donor and acceptor may fall apart. Bright field image shows the unattached LD (red arrow marked) with MΦ. (C). (i) In condition 1, Fl. Intensity of GFP is reduced and an increase in Rhodamine intensity was observed thus signifying the strong FRET between KMP-11 tagged GFP and macrophage membrane bound Rhodamine. (ii). Insignificant change in emission spectra was observed in condition 2.

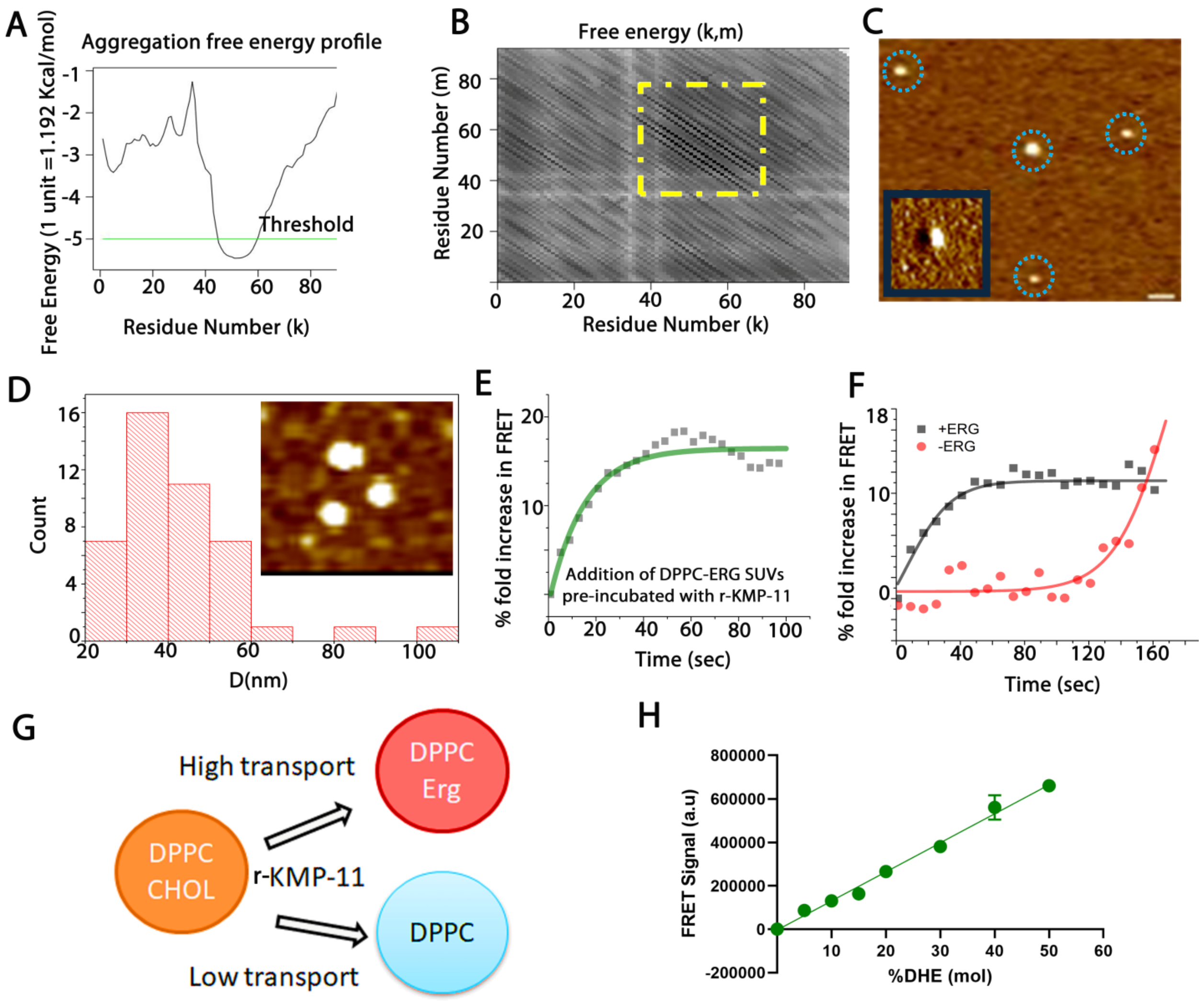


**Figure EV7: Oligomerization of KMP-11.** (A). Using PASTA 2.0 algorithm, we calculated the aggregation/oligomerization free energy that shows 40-60 AA stretch in KMP-11 is prone to oligomer formation. (B). The free energy pairing matrix shows the oligomerization/aggregation prone region (yellow box) in KMP-11 sequence space as evaluated using PASTA 2.0 algorithm (http://protein.bio.unipd.it/pasta2/.) (C). AFM topographic image of r-KMP-11 oligomers formed in presence of membrane. Inset shows the phase topographic image of a single particle. The scale bar is 100 nm. (D). The histogram plot shows the distribution of the oligomeric particles obtained from the AFM images. The inset shows the zoomed in image of the oligomers. CHOL transport property of r-KMP-11. (E). Temporal fold increase in FRET signals. Percentage of fold increase in FRET signal between the transferred DHE (donor) and DAUDA (acceptor) after the mixing of DPPC-DHE SUVs and DPPC-DAUDA-ERG SUVs (which were incubated with r-

KMP-11 for 30 minutes). We performed this experiment to understand the mechanism of cholesterol transport. We incubated r-KMP-11 with DPPC-ERG-DAUDA SUVs assuming the oligomerization in 30 minutes window. Results suggested that the oligomers are particularly responsible for CHOL transport presumably through the hydrophobic tunneling mechanism. (F). Temporal fold increase in FRET signal for DPPC-DHE SUVs and DPPC-DAUDA SUVs (+/- ERG). Our data showed that ERG incorporation in the acceptor vesicles is showing significantly higher CHOL transport rate in comparison to the acceptor vesicles those do not contain ERG. (G). The schematic shows that KMP-11 driven cholesterol transport from DPPC-CHOL membrane is accelerated by the presence of ERG in the acceptor membrane. This data further infers that parasite membrane (that contains ERG) can facilitate the CHOL transport from the host macrophage membrane by KMP-11 during attachment. (H). FRET signal between DHE-DPPC liposome(donor) and DAUDA DPPC liposome (acceptor) in presence of r-KMP-11(20μM)with increasing concentration of DHE(CHOL) in donor liposomes.

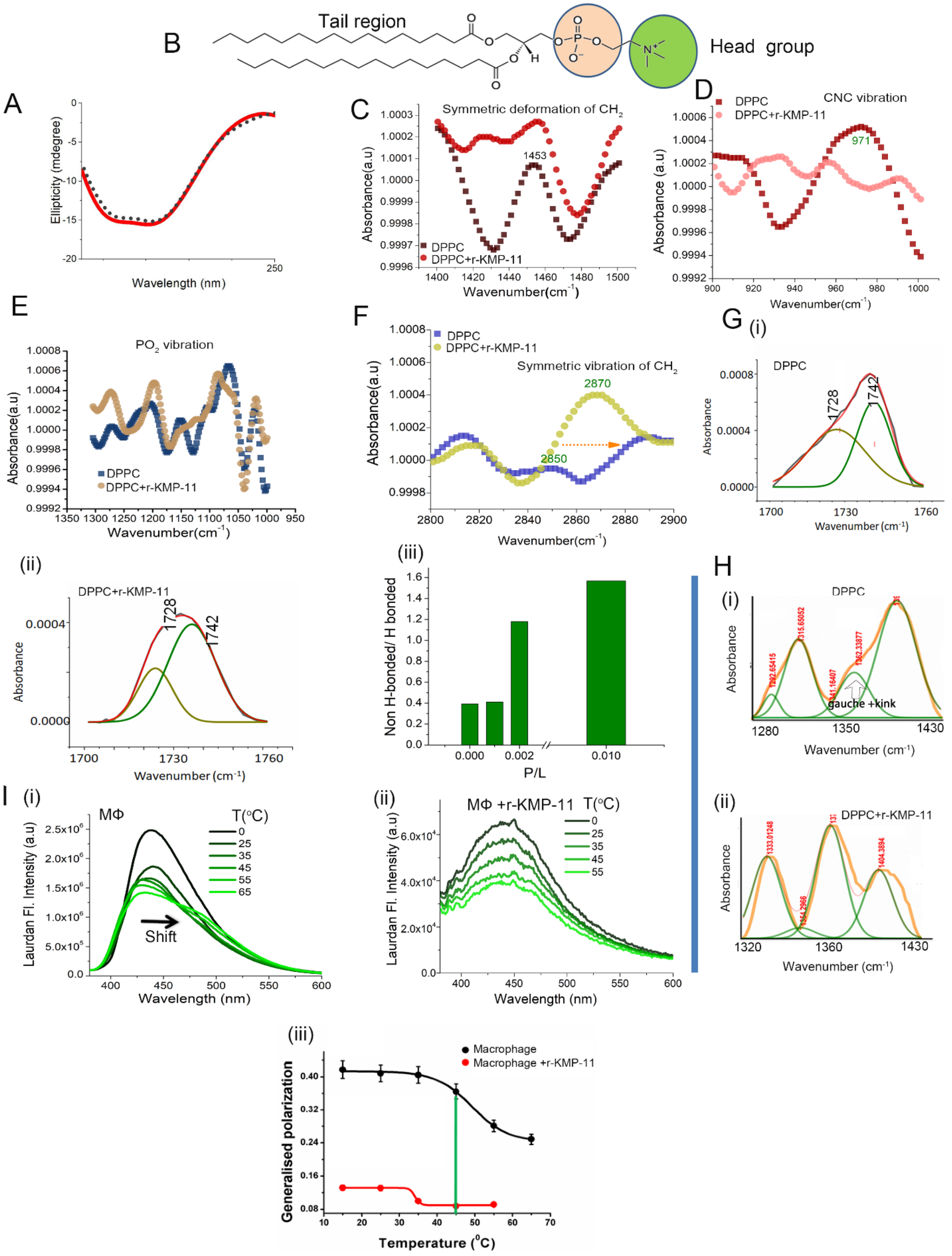
B
Tail region
Head group
A
Ellipticity (mdegree)
Wavelength (nm)
C
Symmetric deformation of $CH_2$
1453
Absorbance(a.u)
Wavenumber($cm^{-1}$)
DPPC
DPPC+r-KMP-11
D
CNC vibration
971
Absorbance (a.u)
Wavenumber($cm^{-1}$)
E
$PO_2$ vibration
Absorbance(a.u)
Wavenumber($cm^{-1}$)
F
Symmetric vibration of $CH_2$
2870
2850
Absorbance(a.u)
Wavenumber($cm^{-1}$)
G (i)
1728
1742
Absorbance
Wavenumber ($cm^{-1}$)
(ii)
(iii)
Non H-bonded/ H bonded
P/L
H
(i)
gauche +kink
Wavenumber ($cm^{-1}$)
(ii)
I (i)
MΦ
T(°C)
Shift
Laurdan Fl. Intensity (a.u)
Wavelength (nm)
(ii)
MΦ +r-KMP-11 T(°C)
(iii)
Macrophage
Macrophage +r-KMP-11
Generalised polarization
Temperature (°C)

**Figure EV8: Interactions induced protein-lipid conformational perturbations.** (A) Far UV-CD spectra of r-KMP-11 in the absence and presence of DPPC membrane. No significant conformational change in the secondary structure of r-KMP-11 due to membrane binding was evident from the CD data. Membrane perturbations study due to KMP-11-lipid binding. (B). The molecular structure of DPPC lipid in which the head group and tail group regions are marked. FTIR signatures of (C) C-N-C, (D). $PO_2$. (E). Symmetric deformation of $CH_2$ (F). Symmetric vibrations of $CH_2$, both in absence and presence of r-KMP-11. (G). Deconvolution of carbonyl (C=O) stretching vibrations in (i) absence and (ii) presence of r-KMP-11 to measure the population of hydrogen bonded carbonyl frequency (appears at 1728 $cm^{-1}$) and nonhydrogen bonded frequency (at 1742$cm^{-1}$). Interestingly, r-KMP-11 binding increased the population of nonhydrogen bonded carbonyl frequency in a concentration dependent manner as shown in (iii). Moreover, enhanced nonhydrogen bonded vibrational states due to r-KMP-11 binding also indicated bilayer thinning. (H). Deconvoluted FTIR spectral signatures of the $CH_2$ wagging band frequency of DPPC in absence (i) and the presence (ii) of r-KMP-11. This FTIR data suggested the increase of gauche rotamers of DPPC due to protein binding. (I). Measurement of MΦ membrane fluidity in the absence and presence of KMP-11. Plot of laurdan emission intensity of labeled MΦ with increasing temperature in the (i). absence and (ii). presence of r-KMP-11. (iii). Generalized polarization (GP) of laurdan labeled macrophage membrane with respect to temperature in absence (black) and presence (red) of r-KMP-11. Typically, concentration of r-KMP-11 was taken 50 μM.

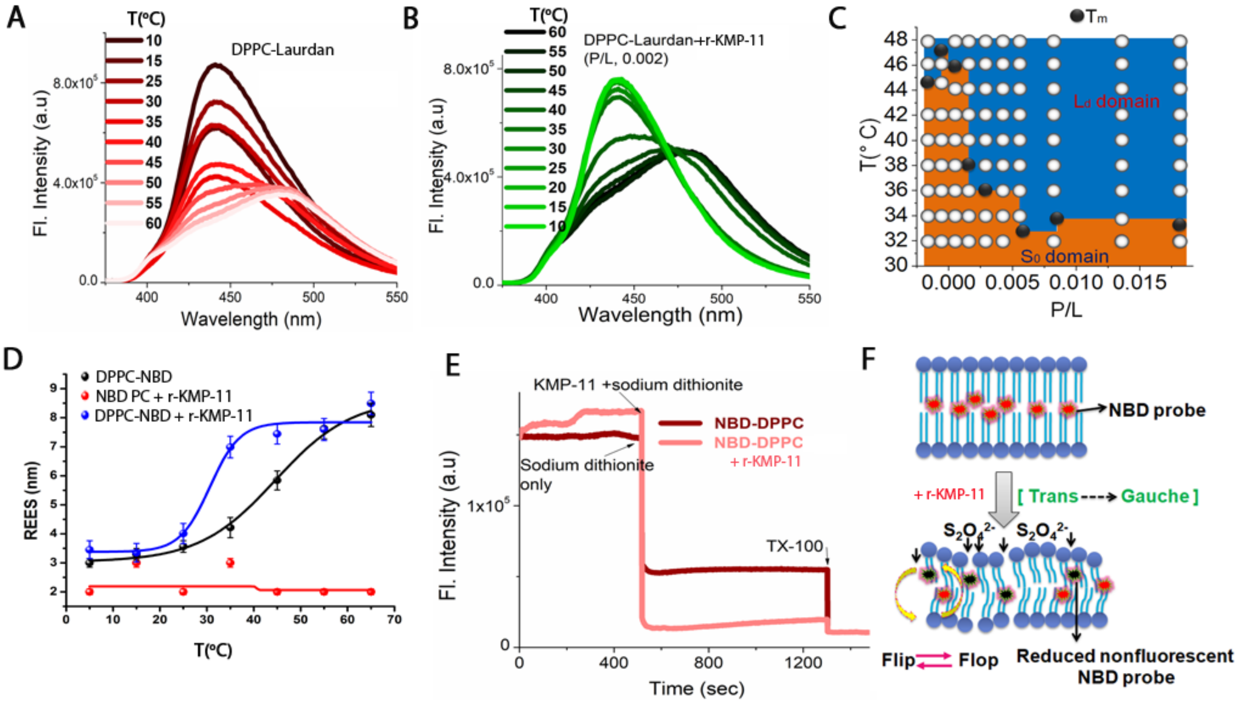


**Figure EV9: Measurements of the governing factors behind KMP-11 induced phase transition.** DPPC laurdan fluorescence in absence and presence of r-KMP-11 with increasing temperature when P/L ratios were kept (A).0, (B). 0.002. The color label indicates the experimental temperatures. (C). Regime diagram illustrating the changes in DPPC phase transition temperatures at different L/P ratios. (D). REES of DPPC-NBD PC and NBD-PC in absence and presence of r-KMP-11. P/L ratio was kept at 0.002 for these experiments. (E). Sodium dithionite assay for measuring the flip-flop dynamics in NBD-PC in presence and absence of KMP-11 which suggested that flip flop motion was significantly increased due to r-KMP-11 binding. (F). Schematic describes that KMP-11 increases the inner flip-flop motion in bilayer as evident from the quenching of NBD which is tagged on the nonpolar side of DPPC. This increased flip flop motion also accompanied trans to gauche conformational turn-over.

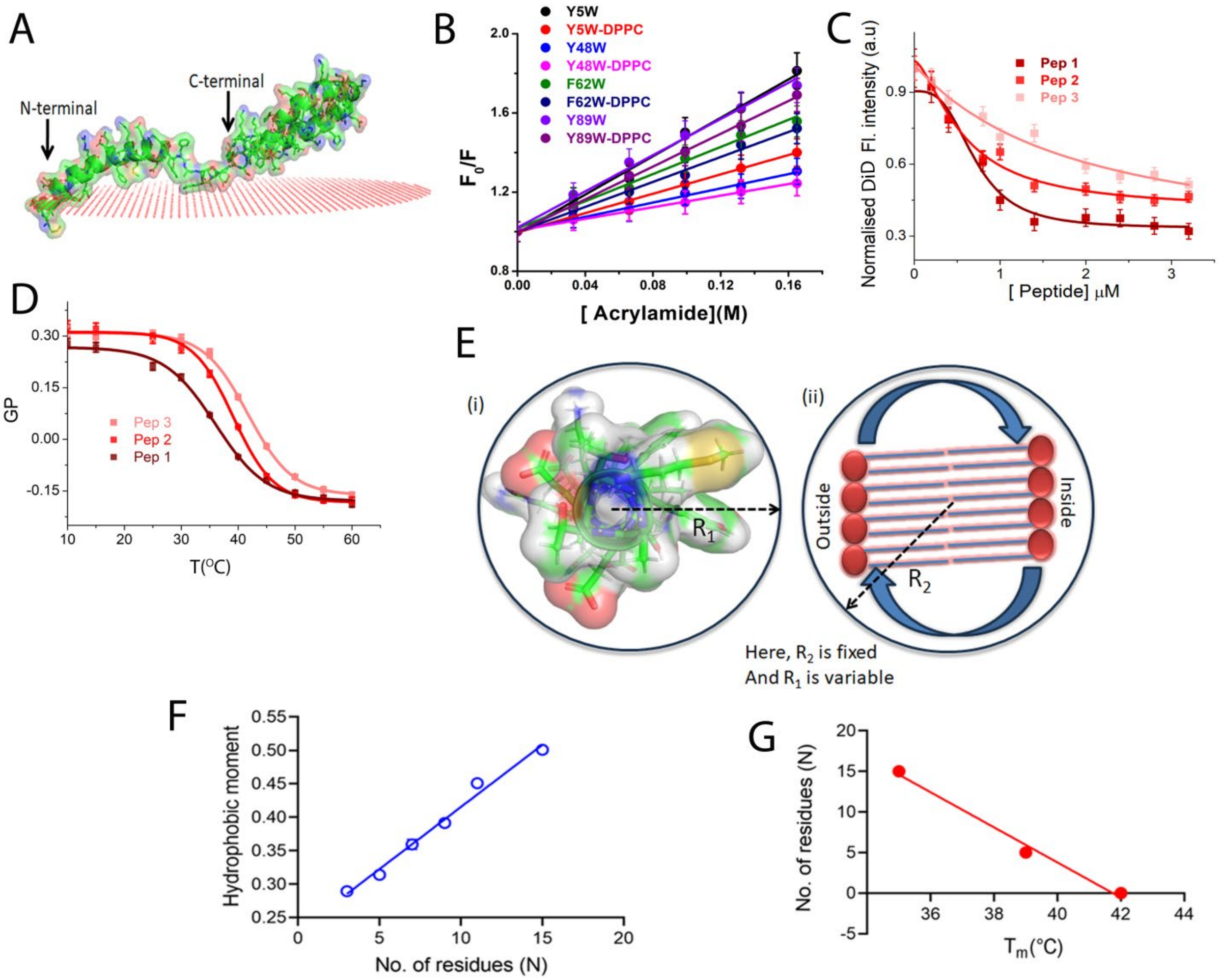


**Figure EV10: Amino-terminal domain is responsible for binding and subsequent changes in membrane.** (A). Orientation of KMP-11 model structure in membrane as obtained from OPM computational study. (B). Representative acrylamide quenching plots ($F_0$/F vs acrylamide concentrations) of four mutants i.e Y5W, Y48W, F62W and Y89W in membrane bound and unbound conditions to examine the domain specific perturbations of the protein through the evaluation of stern-volmer constants from the linear fits of the quenching data. (C). Binding of three different peptides with DPPC model membranes as monitored by the decrease in DiD fluorescence intensity with increasing concentration of added peptides. The solid lines come from the fitting of the data using Hill equation. (D). Laurdan generalized polarization of DPPC membrane in presence of different peptides. Typical peptide concentration was chosen 50 μM. Hydrophobic moment and sequence symmetry model. Two interacting domains i.e. the interacting peptide and lipid bilayer are considered spheres. (E). (i) Model Pymol structure of the amino-terminal stretch of KMP-11 (1-19AA), based on which three peptides were synthesized [peptide 1, peptide2 and peptide3]. Here we assume the peptide domain as a sphere model of radius $R_1$. (ii) On the other hand, considering the flip-flop dynamics of the bilayer entity, we assumed the specific interacting domain of the lipid bilayer as another sphere of radius $R_2$. $R_2$ is fixed since we are not changing any property of bilayer whereas $R_1$is variable due to the change in the sequence arrangement. (F). Relation between hydrophobic moment and the mirror sequence constituting residue number of the membrane interacting stretch. Linear plot of hydrophobic moment ($\mu_H$) against the number of residues constituting the mirror sequences (N) of different combinations based on polarity and hydrophobicity of the amino acid residues in the amino-terminal domain of KMP-11 (Table EV5). (G). Relation between mirror stretch constituting residues and chain melting temperatures of phospholipid model membranes. Linear plot of phase transition temperature ($T_m$) of DPPC against the number of residues constituting the mirror sequences (N) of different combinations based on polarity and hydrophobicity of the amino acid residues. The chain melting temperatures were obtained from the laurdan experiments of DPPC model membranes in presence of three synthetic peptides.
